# Bacterial response to spatial gradients of algal-derived nutrients in a porous microplate

**DOI:** 10.1101/2021.06.23.449330

**Authors:** Hyungseok Kim, Jeffrey A. Kimbrel, Christopher A. Vaiana, Jessica R. Wollard, Xavier Mayali, Cullen R. Buie

## Abstract

Photosynthetic microalgae are responsible for 50% of the global atmospheric CO_2_ fixation into organic matter and hold potential as a renewable bioenergy source. Their metabolic interactions with the surrounding microbial community (the algal microbiome) play critical roles in carbon cycling, but due to methodological limitations, it has been challenging to examine how community is developed by spatial proximity to their algal host. Here we introduce a hydrogel-based porous microplate to co-culture algae and bacteria, where metabolites are constantly exchanged between the microorganisms while maintaining physical separation. In the microplate we found that the diatom *Phaeodactylum tricornutum* accumulated to cell abundances ~20 folds higher than under normal batch conditions due to constant replenishment of nutrients through the hydrogel. We also demonstrate that algal-associated bacteria, both single isolates and complex communities, responded to inorganic nutrients away from their host as well as organic nutrients originating from the algae in a spatially predictable manner. These experimental findings coupled with a mathematical model suggest that host proximity and algal culture growth phase impact bacterial community development in a taxon-specific manner through organic and inorganic nutrient availability. Our novel system presents a useful tool to investigate universal metabolic interactions between microbes in aquatic ecosystems.

## Introduction

Metabolic interactions between microalgae and their associated bacteria, the latter sometimes referred to as the algal microbiome, has been recognized as an important contribution to algal carbon cycling in natural [1] and engineered [2] algal-dominated ecosystems. Heterotrophic bacteria consume up to 50% of the carbon fixed by algae [3], and this mineralization process often leads to exchange of metabolites from bacteria to algae, providing a variety of micronutrients such as trace metals [4], vitamins [5, 6], and phytohormones [7], which can be scarce in nature yet are essential for algal growth. Due to the potential influence of the algal microbiome on algal physiology, much effort has been made to identify algal-associated bacterial community structure and understand how it is related to algal diversity. In particular, it has been revealed that the algal-associated bacterial community can be highly conserved across time [8, 9], uniquely shaped by their algal host [10–12] and exhibits structural differences between algal-attached and free-living [12]. These studies indicate that the community-level bacterial response to algal hosts depends at least partially on physical proximity, including physical attachment, as well as the type of algal-excreted products. Indeed, both direct cell-to-cell attachment [10, 11, 13, 14] and chemotaxis towards algal exudates [15–18] are commonly observed phenomena.

A ubiquitous mechanism that explains how heterotrophic bacteria respond to algal-derived organic matter is diffusive transport of metabolites. Metabolites diffusing around an alga create a spatial gradient of nutrients, and the region where the algal-derived organic nutrients are abundant is termed the phycosphere [1, 18, 19]. For an alga sized less than 100 μm, diffusion within its phycosphere plays a major role in transporting metabolites rather than fluid advection [1] or turbulence [20]. Moreover, diffusive transport is considered to be 50 times more efficient for encountering molecules than cellular swimming especially at a low stirring number (defined as *lv* / *D*; *l* is intermolecular spacing, *v* swimming speed and *D* diffusion coefficient) [21, 22]. Therefore, by maintaining spatial proximity to the algal cell inside the phycosphere, bacteria can best exploit algal-derived organic matter through diffusive effects.

In order to assess how microorganisms respond to the diffusion of metabolites such as algal exudates, *in vivo* co-culture methods have been developed. For example, a dual-chamber system separated by a porous membrane [17, 23–25] has enabled further downstream biological analyses, such as plate reader-compatible *in situ* growth measurements [24], metabolomics [23], or microscopy to track cellular motility [17] and biofilm formation [25]. Microscopy has indeed provided valuable information on how bacteria can temporally respond to the presence of algae via chemotaxis [16, 17]. Nonetheless, existing techniques have not yet enabled the co-cultivation of microalgae with multiple bacterial species, especially for time periods long enough to detect community level dynamics (days to weeks).

In this paper we present a porous microplate targeted to the study of *in vivo* interactions between algae and bacteria. By using the microplate we develop a protocol to culture the microorganisms for up to several weeks, overcoming previous limitations mentioned above. We employ the hydrogel poly(2-hydroxethyl methacrylate-*co*-ethylene glycol dimethacrylate) (HEMA−EDMA), previously shown to be an effective building co-polymer to enable cell-to-cell communication ranging from microbial [26] to mammalian cells [27]. By using the porous microplate, we set our central goal to understand how metabolite diffusion plays a role in the interaction between algal cells and their associated bacteria. We explore how bacterial populations respond to algal exudate diffusion by altering its input to bacterial communities through changing the physical proximity between the source (the algae) and the sink (the bacteria). Specifically, by taking advantage of our incubation system, we hypothesize that inorganic nutrients from the medium are converted into dissolved organic carbon (DOC) through algal photosynthetic activity *in vivo*, and that the conversion creates a spatial gradient of DOC and inorganic nutrients across the culture wells, the response to which can be detected by bacterial abundance and community structure changes.

To test our hypotheses, we first examined the model alga *Phaeodactylum tricornutum* grown in the porous microplate where culture wells are spatially arranged to generate different nutritional availability. Then we co-cultured *P. tricornutum* with two commonly algal-associated bacterial strains (genera *Algoriphagus* and *Marinobacter*), where the bacteria were incubated at different distances from the alga. The strains were previously isolated from *P. tricornutum* mesocosms [10, 13] and have been previously shown to affect algal growth [28]. Finally, we incubated *P. tricornutum* in the porous microplate with mixed bacterial communities to observe how community structure responds to physical distance from the algal host.

## Materials and Methods

### Acrylic and polydimethylsiloxane (PDMS) molds preparation

The incubating device for porous microplate was designed using a CAD software (Solidworks, Dassault Systèmes) and the exported drawing files were used to laser cut 1/4” and 1/8” acrylic sheet (Universal Laser Systems; Supplementary Fig. S1). After washing the cut acrylic parts with deionized water, they were attached by acrylic (Weld-On) and epoxy (3M) adhesives which was followed by curing process for ~18 h. Polydimethylsiloxane (PDMS) (Sylgard 184, Dow Corning) was cast onto the acrylic mold and cured at 80°C for at least 3 h. The PDMS mold was carefully detached from the acrylic surface by dispensing isopropyl alcohol (VWR) into the area between the PDMS and the acrylic molds.

### Porous microplate preparation

Synthesis of co-polymer HEMA–EDMA was based on previously described protocols [26, 27] and details are given as follows. Prepolymer solution HEMA−EDMA was prepared by mixing 2-hydroxyethyl methacrylate (HEMA; monomer, 24 wt.%, Sigma-Aldrich), ethylene glycol dimethacrylate (EDMA; crosslinker, 16 wt.%, Sigma-Aldrich), 1-decanol (porogen, 12 wt.%, Sigma-Aldrich), cyclohexanol (porogen, 48 wt.%, Sigma-Aldrich) and 2,2-dimethoxy-2-phenylacetophenone (DMPAP; photoinitiator, 1 wt.%). The solution was stored at room temperature without light exposure until further use. Glass slides (75 × 50 mm^2^, VWR) were chemically cleaned by sequentially soaking in 1 M hydrochloric acid and 1 M sodium hydroxide for one hour, followed by rinsing with deionized water and air drying. The prepolymer solution was cast onto the PDMS mold and a glass slide was placed on the mold. The solution was then polymerized under ultraviolet light with a wavelength 365 nm for 15 min by using a commercial UV lamp (VWR). The photopolymerized device was detached from the PDMS mold and stored in a jar containing methanol (VWR) until further use. The jar was refilled with new methanol twice in order to remove remaining porogen and uncrosslinked monomers from the hydrogel.

Before incubating microbial cells in porous microplate, each device was decontaminated by replacing the solvent with 70% alcohol (VWR) and storing for 24 h. They were immersed in a pre-autoclaved jar with f/2-Si medium for two weeks, where the jar was refilled once with new sterile medium to adjust its pH for algal culture and remove any solvent remaining in the hydrogel.

### Scanning electron microscopy (SEM)

A sample of polymerized HEMA−EDMA was taken out from a containing methanol and the remaining solvent was removed by air drying for at least 1 week. The sample was attached to a stub, loaded on MERLIN^TM^ SEM and imaged with SmartSEM software (Zeiss) with 16,270 times of magnification at the Electron Microscopy Facility in the Massachusetts Institute of Technology Materials Research Science and Engineering Centers (MIT MRSEC).

### Strains and culturing conditions

Axenic *P. tricornutum* CCMP 2561 was acquired from the National Center for Marine Algae and Microbiota (NCMA). *P. tricornutum* was maintained in f/2-Si medium with 20 g L^−1^ commercially available sea salts (Instant Ocean, Blacksburg) [10, 13]. Batch cultures were grown at 20°C with a 12 h light/12 h dark diurnal cycle and a light intensity of 200 μmol photons m^−2^ s^−1^ (Exlenvce). Every two weeks, axenic cultures were monitored for bacterial contamination by streaking culture samples on marine broth agar [29]. Growth of *P. tricornutum* was measured by counting cells using a hemocytometer or flow cytometry. Specific growth rates were calculated from the natural log of the cell densities in triplicate at the exponential growth phase (day 3 for the batch culture, day 5 for the porous microplate system).

Bacterial community samples (referred to as “phycosphere enrichments”) were obtained from mesocosms of *P. tricornutum* and maintained as described by Samo *et al*. [13]. Two bacterial strains, *Marinobacter* sp. 3-2 and *Algoriphagus* sp. ARW1R1, were isolated from phycosphere enrichment samples (Supplementary Table S2). The isolates were either maintained by growing on marine broth agar plates at 30°C or by co-culturing with *P. tricornutum* through inoculation of a single colony into the axenic culture.

### Single bacterial isolates experiment

*Marinobacter* sp. 3-2 and *Algoriphagus* sp. ARW1R1 were prepared by inoculating a single colony into marine broth and growing overnight (30°C, 150 r.p.m.). The cells were washed with f/2-Si twice by centrifuging at 2,258 rcf for 4 min, diluted to a density of ~4 × 10^6^ cells ml^−1^ (*Marinobacter* sp. 3-2) and ~4 × 10^5^ cells ml^−1^ (*Algoriphagus* sp. ARW1R1). Diluted cells were then inoculated into the microplate wells. The cell densities were initially set to resemble *in situ* conditions with *P. tricornutum*-associated bacterial communities where *Marinobacter* displayed relative abundances several folds higher than *Algoriphagus* [13]. *P. tricornutum* was acclimated to the microplate environment by inoculating the stationary phase-culture in a separate microplate. After incubating for 4 d, ~1 × 10^7^ algal cells ml^−1^ were transferred to the center well of the experimental device with added bacteria in the surrounding wells. Three replicated microplates were placed in a single transparent covered container (128 × 85 × 10 mm^3^, VWR) which was filled with ~25 ml f/2-Si medium to keep the devices hydrated throughout the incubation period of 20 d. The procedures were conducted under a biosafety cabinet to prevent any biological contamination.

Every 2–3 days, 5 μl samples were collected from wells and transferred to a 96-well plate (Corning). Cells were then diluted 35 times with 1x phosphate buffered saline solution (PBS) (Mallinckrodt), followed by addition of 16% formaldehyde (Thermo Scientific) to a final concentration of 2% adjusting to a final volume of 200 μl. Fixed samples were stored at 4°C for no more than 4 weeks. After collecting the cell samples for 20 d, the remaining cultures were taken out from the device and were streaked on marine broth agar plates to check if any of the culture wells had been cross-contaminated. Five out of 108 bacterial samples were cross-contaminated, and they were excluded from the analysis. No less than three replicates were retained after excluding the cross-contaminated samples.

### Flow cytometry

Fluorescent counting beads for flow cytometry (Alignflow^TM^, Thermofisher) were diluted 20 times with PBS and 50 μl was added to each well of the 96-well plate for calibration. SYBR Green I nucleic acid stain (Thermofisher) was added to the samples with a final concentration of 0.1% v/v, and allowed to sit at room temperature at least for 30 min without light exposure. Flow cytometry analysis was conducted on a BD FACS Canto II HTS to quantify the number of algal and bacterial cells with parameter setting as follows: (voltage) FSC = 580, SSC = 370, GFP = 400, PE = 330, PerCP = 647, PE-Cy7 = 677, Alexa Fluor 680 = 290, APC-Cy7 = 410, Pacific Blue = 440, AmCyan = 539. Populations were plotted with GFP-A and APC-Cy7-A intensities, allowing a clear distinction between counting beads, stained algal and bacterial cells. The data were exported to .csv files using FlowJo (BD) and converted into cell density data using MATLAB (Mathworks).

### Bacterial community experiment

To examine bacterial community structure changes induced by the presence of the diatom, ~1 × 10^6^ *P. tricornutum* cells ml^−1^ were placed in the center well and the results were compared to control incubations where sterile media was placed in the center. Phycosphere enrichment samples [10, 13] (with the algal cells removed with a 0.8 μm filter) were used as a bacterial community inoculant where 100 μl samples were placed in the surrounding wells in the hexagonal array. On the outmost well, 100 μl sterile f/2-Si medium was inoculated as a blank control for 16S rRNA community analysis.

The devices were incubated in a container (GasPak™ EZ container systems, BD) which was filled with f/2-Si medium. All preparation steps were performed in a laminar flow hood to prevent any bacterial contamination. The container was incubated under the same condition as described above. After 7 d, each well sample was collected and filtered into a 96-well filter plates (0.2 μm pore size, Pall AcroPrep) to remove liquid, and was stored in −20°C until further analysis.

### 16S ribosomal RNA gene amplification and sequencing

Twenty microliters of sterile DNA-free water was added to each well of the filter plates, and cells were lysed to release the DNA with heat (95°C for 10 minutes). The 16S rRNA gene was directly amplified without cleanup, using primers targeting the v4 region and modified to include Illumina platform adaptor sequence on the 5’ ends. The PCR contained 10 μl of 5 Prime MasterMix, 1 μl of 10 μM of forward 16S primer (5’- TCGTCGGCAGCGTCAGATGTGTATAAGAGACAGGTGYCAGCMGCCGCGGTAA-3’), 1 ul of reverse 16S primer (5’- GTCTCGTGGGCTCGGAGATGTGTATAAGAGACAGGGACTACNVGGGTWTCTAAT – 3’), and 5 μl of DNA template. Cycling conditions were as follows: denaturation at 94°C for 3 min, followed by 30 cycles of denaturation at 94°C for 45 s, annealing at 51°C for 30 s and extension at 72°C for 1.0 min. The final extension was conducted at 72°C for 10 min and the samples were held at 4°C. A second round of amplification added Illumina Dual Nextera XT indexes and sequencing adaptors [30]. Each library was quantified with a Qubit broad range dsDNA assay and equimolar amounts of each library were pooled. The size and concentration of the final pool was verified by a D5000 High Sensitivity assay on the Agilent Tapestation. Six pM of the pooled libraries were combined with 15% phiX and paired end sequenced on an Illumina MiSeq for 500 cycles.

Paired-end MiSeq reads were filtered to remove read pairs that contained Illumina adapter or barcode kmers with bbduk v38.22 using options k=31 and hdist=1 [31]. Read trimming and quality filtering was done with DADA2 v1.12.1 using options trimLeft = c(4, 14), truncLen = c(200,150), maxN = 0, maxEE = c(2, 2), truncQ = 2 [32]. Read pairs remaining after filtering were further processed into amplicon sequence variants (ASVs) retaining ASVs of length 274 nucleotides with at least 2 reads in at least 2 samples. The sequences were then aligned with MUSCLE v3.8.1551 [33] and clustered into an approximately-maximum-likelihood tree with Fasttree v2.1.10 [34]. Taxonomy was assigned using IDTAXA [35] against the Silva version 132 SSU database [36]. ASVs were analyzed using the R package phyloseq v1.30.0 [37]. Samples were quality-filtered by removing taxa that were seen less than 10 times in 3 samples and normalized by Cumulative Sum Scaling (CSS) method using the R package metagenomeSeq v1.28.2 [38]. In brief, a mean of 95,770 (from 11,535 to 241,656) and 19,997 (from 61 to 159,617) read pairs were sequenced per sample of bacterial wells and of negative controls, respectively (Supplementary Table S3). Among a total of 120 samples, 12 negative controls were used to validate the sequencing results and were not statistically analyzed further. Another 36 samples intermediate to layers 1 and 2 were not assigned with a layer number and were excluded from the statistical analysis. R package phyloseq v1.30.0 was used to perform principle coordinate analysis (PCoA), vegan v2.5.6 to perform permutational multivariate analysis of variance (PERMANOVA) [39] based on weighted Unifrac [40], and R function (aov) was used to perform one-way and two-way analysis of variance (ANOVA) tests using packages tidyverse v1.3.0 [41], vegan v2.5.6 [42].

## Results

### *P. tricornutum* accumulates in a porous microplate

HEMA–EDMA is a nanoporous hydrogel and can easily dry out in standard laboratory conditions (Fig. 1a,b). To circumvent the dehydrating issue, hydrogel microplates were immersed in f/2-Si medium in a sealed container where ~95% of the initial 100 μl volume was retained over 2 weeks (Fig. 1c, Supplementary Table S1). We measured cell abundances of *P. tricornutum* incubated in the microplate and compared them with conventional batch cultures (Fig. 2a). Unexpectedly, *P. tricornutum* in the porous microplate reached a density of (2.3 ± 0.8) × 10^8^ cells ml^−1^, which was ~20 folds higher than those under a batch condition (Fig. 2b). However, the two culture methods did not show a difference in the growth rate, suggesting the hydrogel did not impact *P. tricornutum* physiology (*P* = 0.413, Fig. 2b inset).

**Figure 1.**
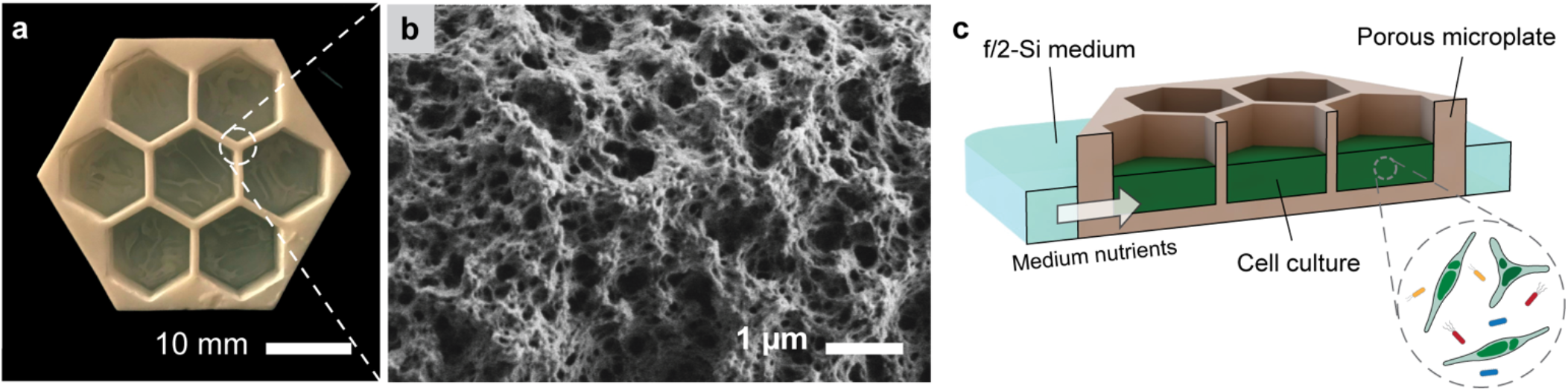
Porous microplate for nutrient diffusion. (a) Device with a hexagonal culture well array (wall thickness: 0.9 mm). (b) Scanning electron microscopy image of nanoporous HEMA–EDMA. (c) Cross-sectional schematic illustrating microbial cell incubation in a porous microplate.

**Figure 2.**
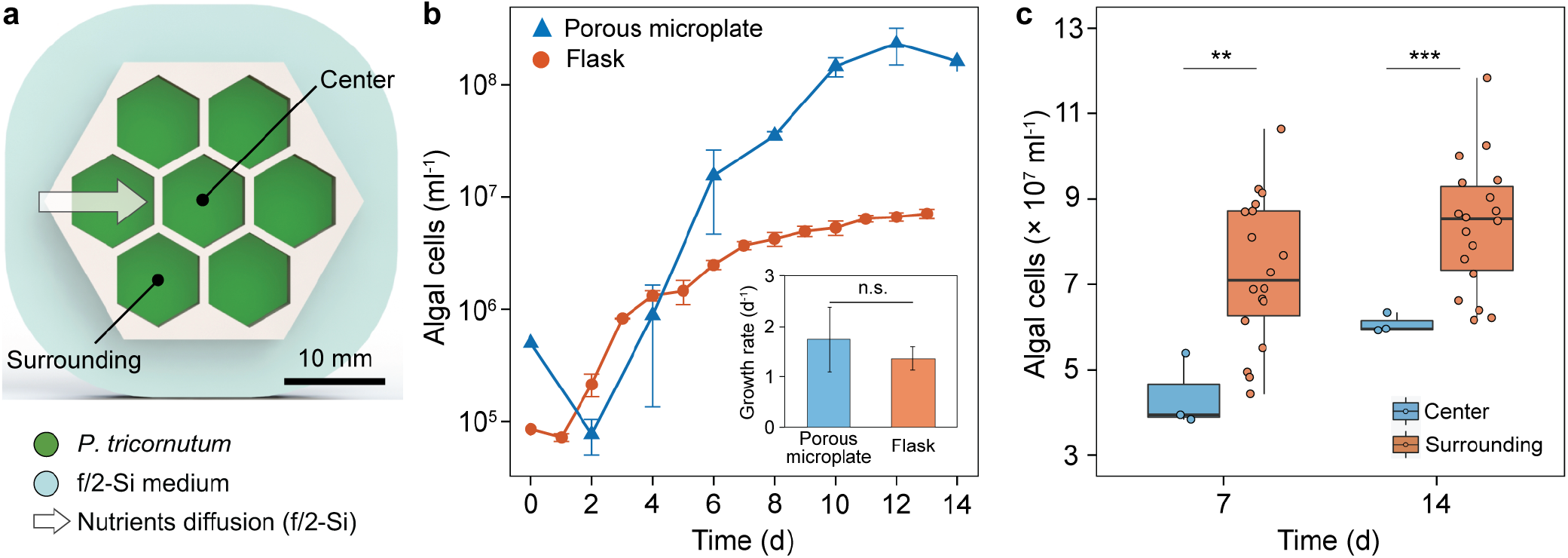
Cultivation of *P. tricornutum* in the porous microplate. (a) Schematic of a microplate for algal culture. (b) Growth curve and maximal growth rate (inset) comparing the porous microplate with flask culture. Error bars, standard deviation of triplicates. (c) Cell abundance at center (*n* = 3) and surrounding (*n* = 18) wells after incubation. Asterisks denote statistical differences with following levels (two-tailed *t*-test): *** (*P* < 0.001), ** (*P* < 0.01), * (*P* < 0.05) and n.s. (not significant).

We hypothesized that the higher carrying capacity in the microplate might be caused by the f/2-Si medium nutrients diffusing into the well as the algae grew, since nutrients are constantly consumed and a concentration gradient is created across the porous HEMA–EDMA [26]. To test this hypothesis, we designed a hexagonal array with one center and six outer wells, so that nutrients outside the microplate can supplement the outer wells faster than the inner wells (Fig. 2a). Indeed, algal populations cultured in the outer wells reached higher concentrations than cultures in the center (*P* < 0.01, Fig. 2c), but those at both locations still exhibited higher abundances than standard batch cultures. One further experiment confirmed that the inorganic nutrients in the outer reservoir diffused to the culture wells; we compared *P. tricornutum* abundances in unaltered medium with post-experiment reservoir medium, testing if the spent medium had lower nutrient concentrations. As expected, *P. tricornutum* grew slower and to lower concentrations in this spent medium compared to fresh medium, consistent with the hypothesis of inorganic nutrient diffusion from the reservoir into the wells (Supplementary Fig. S2).

### Bacteria respond to gradients of *P. tricornutum* exudate and medium nutrients

We next explored how algal-associated bacteria grow and respond to the diffusion of algal and medium nutrients in the porous microplate. In a hexagonal array of microplate wells, *P. tricornutum* and two bacterial strains, *Marinobacter* sp. 3-2 and *Algoriphagus* sp. ARW1R1, were co-cultured at different distances from one another (Fig. 3a). The two bacterial strains were previously isolated from *P. tricornutum* enrichments [13] and are relatively abundant in laboratory cultures where bacterial growth depends on *P. tricornutum* photosynthate [10]. The abundances of the two strains at different time points were compared to one another as a function of the distance away from the center well, which contained either *P. tricornutum* or f/2-Si medium.

**Figure 3.**
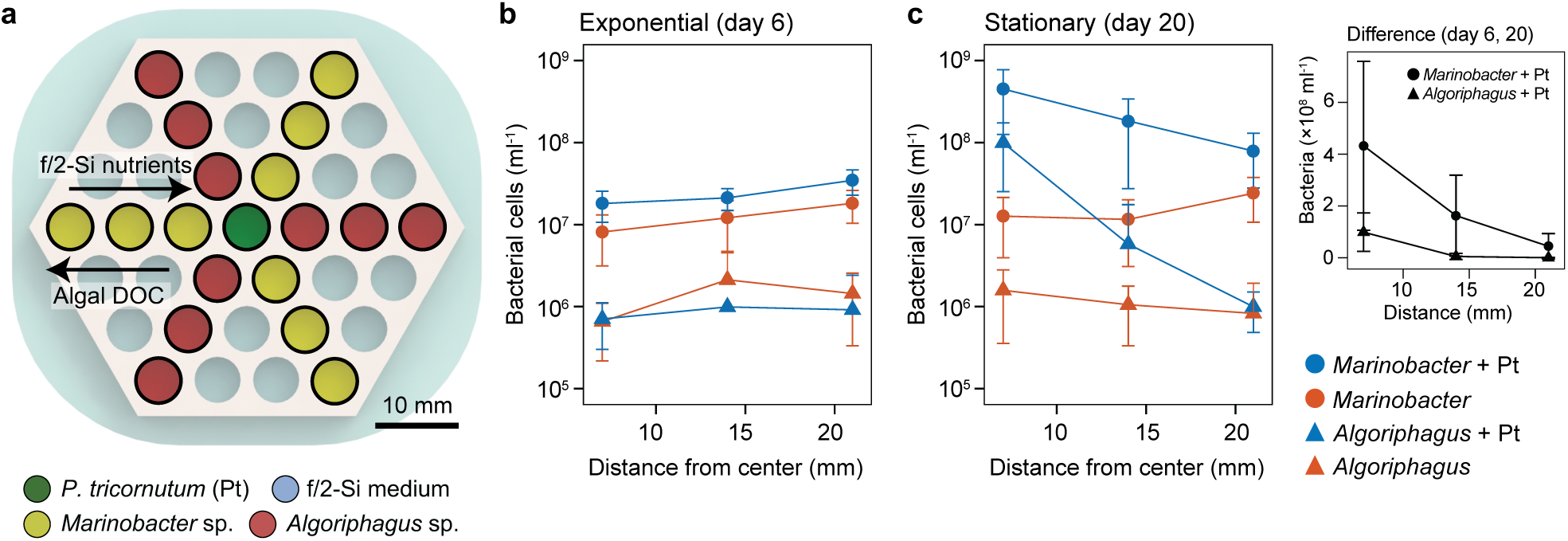
Measurements of bacterial growth in the porous microplate. (a) Schematic of a microplate depicting well locations of *P. tricornutum* and two bacterial strains, *Marinobacter* sp. 3-2 and *Algoriphagus* sp. ARW1R1. Abundance of the two bacterial strains sampled at (b) exponential and (c) stationary phase; (inset) difference of the abundance between the timepoints. Error bars, standard deviations of the following number of replicates: *n* = 5 (*Algoriphagus* with Pt, distance 21 mm), *n* = 8 (*Algoriphagus* without Pt, distance 21 mm), *n* = 9 (all others).

During the initial algal growth phase (day 0–6), we observed a rapid increase of bacterial abundances (Supplementary Fig. S3). Interestingly, *Marinobacter* was more abundant as distance from the center well increased, regardless of the presence of *P. tricornutum* (*P* < 0.05, Fig. 3b). However, *Algoriphagus* did not exhibit any such difference (*P* = 0.814 with algae and *P* = 0.165 without). This tendency of higher abundances in the outer wells was also observed for *Marinobacter* without *P. tricornutum* at the other timepoints in the early growth phase (day 2–8, *P* < 0.05, Supplementary Fig. S3).

We expect that medium inorganic nutrients were consumed by algal photosynthesis, as shown in our previous experiments, creating a concentration gradient of these inorganic nutrients increasing towards the outer microplate wells. This suggests that *Marinobacter* sp. 3-2 growth depended on the inorganic nutrients as well as algal DOC at this early growth phase. For the treatment without *P. tricornutum* where we also detected increasing growth towards the outer wells, there was likely also a gradient in inorganic nutrient concentration. This is because the outer wells are adjacent to the medium input outside the microplate, whereas the inner wells are in the center and adjacent to other wells with potentially competing bacteria. Nonetheless, *Marinobacter* co-cultured with algae were more abundant than without (*P* < 0.001, Fig. 3b), suggesting that the bacteria also used some metabolites from *P. tricornutum* for growth and that these metabolites were able to diffuse all the way to the outermost layer.

Results from later in the growth phase (e.g. day 17–20) were drastically different, where the two bacterial strains exhibited distinct responses to the spatial arrangement. On day 20, the abundances of both strains strongly increased with proximity to *P. tricornutum*, indicating that their growth was correlated to algal DOC diffusion (*P* < 0.001, Fig. 3c). *Algoriphagus* appeared to have a greater dependence on *P. tricornutum* exudates than *Marinobacter*, as measured by a stronger decrease in its abundance with increasing distance from the center (*Algoriphagus*, 97-fold; *Marinobacter*, 6-fold). Furthermore, in the outermost well, *Marinobacter* was 100 times more abundant than *Algoriphagus*, suggesting that *Marinobacter* does not require spatial proximity to *P. tricornutum* for its growth (note that initial *Marinobacter* abundance was ten times greater than *Algoriphagus*). Indeed, the growth of *Algoriphagus* located further away from *P. tricornutum* (in the outer wells) was negligible over the 20 day incubation period, whereas *Marinobacter* grew in all the wells (Supplementary Fig. S3). The innermost well was the only location where *Algoriphagus* grew, suggesting the algal DOC needed by *Algoriphagus* did not diffuse further, or was fully incorporated by *Algoriphagus* cells in the innermost well.

### Spatial influence on bacterial community development

Our previous experiment documents the differential response of two bacterial strains grown in monoculture exchanging metabolites with the algal host and potentially with one another. The next step was to test our system with mixed communities and examine how community development is influenced by distance from the alga. By inoculating *P. tricornutum* in the center well and by filling the container with f/2-Si medium outside (Fig. 4a), we expect that a spatial gradient of increasing inorganic nutrients (higher outside, lower inside) and an opposite gradient of algal DOC (higher inside, lower outside) was generated along the well array, similar to gradients that are hypothesized to exist in the phycosphere at the single cell level. The algal and bacterial cells were incubated in the microplate for a week, a period long enough to elicit population level changes in previous experiments, as discussed above. Bacterial communities were then analyzed by sequencing their 16S ribosomal RNA (rRNA) genes, enabling us to examine how community development is affected by physical location along the two gradients.

**Figure 4.**
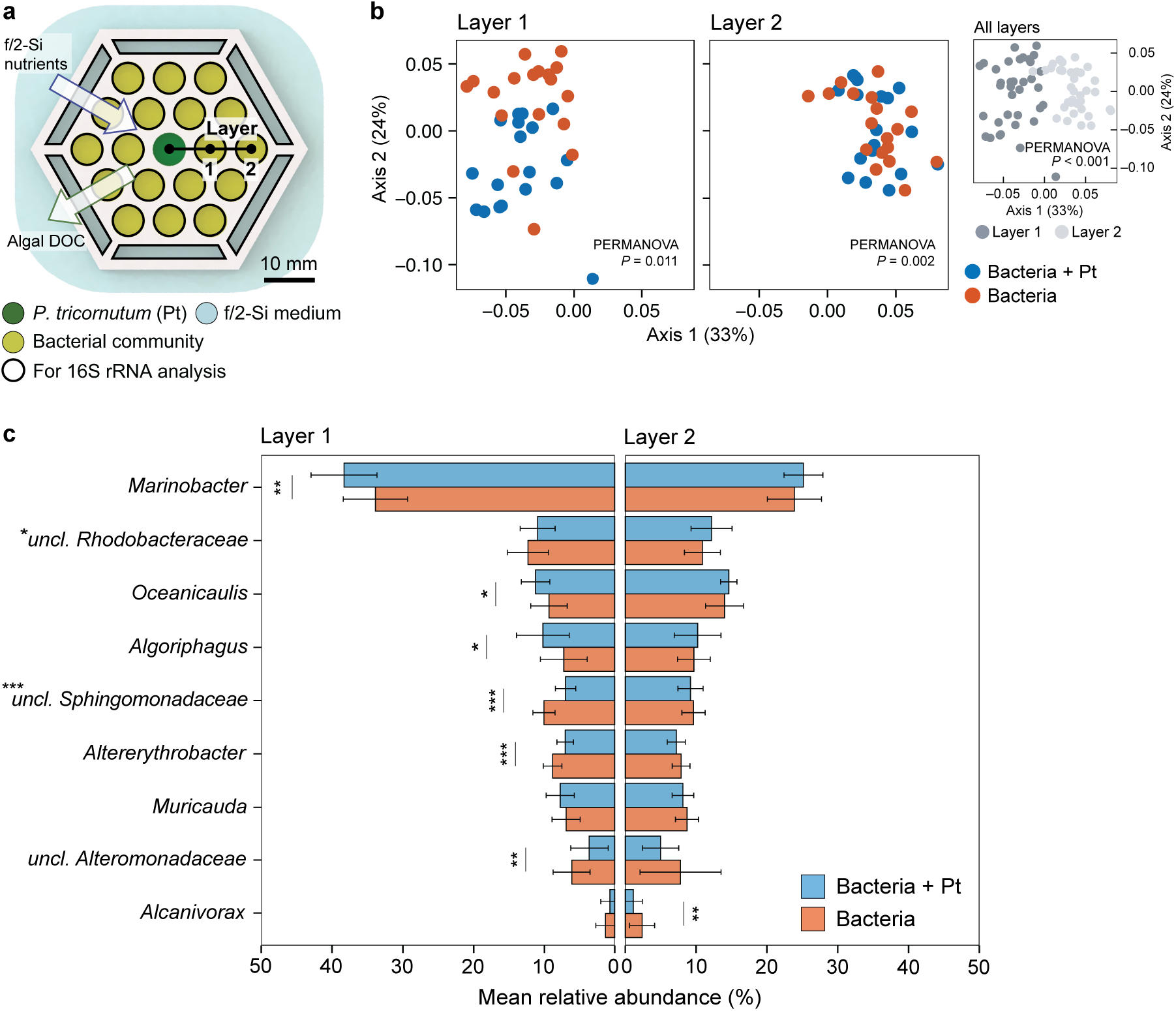
Bacterial community analysis in the porous microplate. (a) Schematic of a microplate depicting locations of *P. tricornutum* and associated bacterial communities. (b) Principle coordinate analysis of the communities with *P*-values (PERMANOVA) on structural difference between layer conditions (inset) and the two treatments. (c) Genus level taxonomic abundance of the communities comparing two treatments incubated for 7 days (means more than 0.5% displayed). Error bars, standard deviations of 18 replicates. Asterisks next to genera (y-labels), statistically significant interaction term of layer number and treatment (two-way ANOVA). Asterisks next to color bars, statistical significance of differences between two treatments (one-way ANOVA) with following levels: *** *P* < 0.001, ** *P* < 0.01, * *P* < 0.05.

For a global analysis, samples were grouped by treatment (with or without *P. tricornutum* in the middle well) and layer number (layer 1 located at the inner and layer 2 at the outer well). Data visualized using principal coordinate analysis (PCoA) using weighted Unifrac metric [40] showed a notable difference between layer 1 and layer 2, regardless of the treatment (*P* < 0.001, Fig. 4b inset). In both layers, the sample groups were statistically different between treatments (*P* < 0.05), but the structural difference was most visibly obvious in layer 1 compared to layer 2 with a higher proportion of the variance explained by the principal coordinates (Fig. 4b, Supplementary Fig. S4). This suggests that the community structure was indeed affected by algal photosynthetic activity and exudate production, and to a greater extent for bacterial cells located closest to *P. tricornutum*.

To investigate in greater detail which taxa were influenced by spatial proximity to the algal cells, relative abundances were quantified by merging amplicon sequence variant (ASV) reads at the genus level and compared between well location (inner vs. outer) and treatment (with and without algae in the center well). Compared between treatments, the communities incubated closest to the center well (layer 1) contained more genera that were statistically different in their abundance than in layer 2 (Fig. 4c). In detail, six genera, *Marinobacter*, *Oceanicaulis*, *Algoriphagus*, *unclassified Sphingomonadaceae*, *Altererythrobacter* and *unclassified Alteromonadaceae*, were different in layer 1, whereas only *Alcanivorax* showed such difference in layer 2 (*P* < 0.05). We also observed an interaction between treatment and well location with genera *Marinobacter*, *unclassified Rhodobacteraceae*, *unclassified Sphingomonadaceae*, *Muricauda* and *Altererythrobacter,* suggesting that they were affected by physical proximity to *P. tricornutum* in the center well and inorganic nutrients on the outside of the microplate (*P* < 0.05). Specifically, the genera *Marinobacter*, *Oceanicaulis*, *Algoriphagus*, and *Muricauda* exhibited higher relative abundances in co-cultures with algae compared to without, suggesting that their response to the algal exudate was stronger than their response to inorganic nutrients. On the other hand, the genera *unclassified Rhodobacteraceae*, *unclassified Sphingomonadaceae*, *Altererythrobacter* and *unclassified Alteromonadaceae* were relatively less abundant with algae compared to without, indicating they were more strongly influenced by inorganic nutrients, or more likely did not respond to algal exudates as well as the other taxa.

### Predictive modeling of nutrient concentrations around an alga

The presumed differential bacterial assimilation of inorganic nutrients and algal-derived organic matter in our porous microplate provides a testable analogous model to the microscale processes that likely occur at the single cell level in the phycosphere [19]. Our system did not specifically test algal-attached bacteria, but the location of the wells at different distances from the algal culture enabled us to artificially create different zones comprised of different concentrations of both algal-derived metabolites and inorganic nutrients. Similarly, at the scale of the single algal cell, its photosynthetic activity creates two contrasting concentration gradients governed by the laws of diffusion: an increase of algal DOC and a decrease of inorganic nutrients towards the alga (Fig. 5a). Furthermore, the concentration profiles will also be affected by culture aging as cell-to-cell distances and the types of released compounds change [19, 43–47]. With these assumptions, we calculated spatial concentrations of algal DOC and inorganic nutrients at the scale of a cell and under different growth phases (Supplementary Note S1). We found algal released compounds decreased by 3–7-fold with increasing distance from the cell surface, supporting the theoretical validity of the phycosphere [1, 19] regardless of growth phase (Fig. 5b). However, as inorganic nutrients are rapidly consumed during culture growth [45, 48], its overall concentrations become very low when it reaches stationary phase, and this is unaffected by closeness to the algal cell. This is represented by a drastic difference in the spatial concentrations of nitrate (as a proxy of inorganic nutrients) between early (day 0–6) and late growth phases (day 6–14, Fig. 5c).

**Figure 5.**
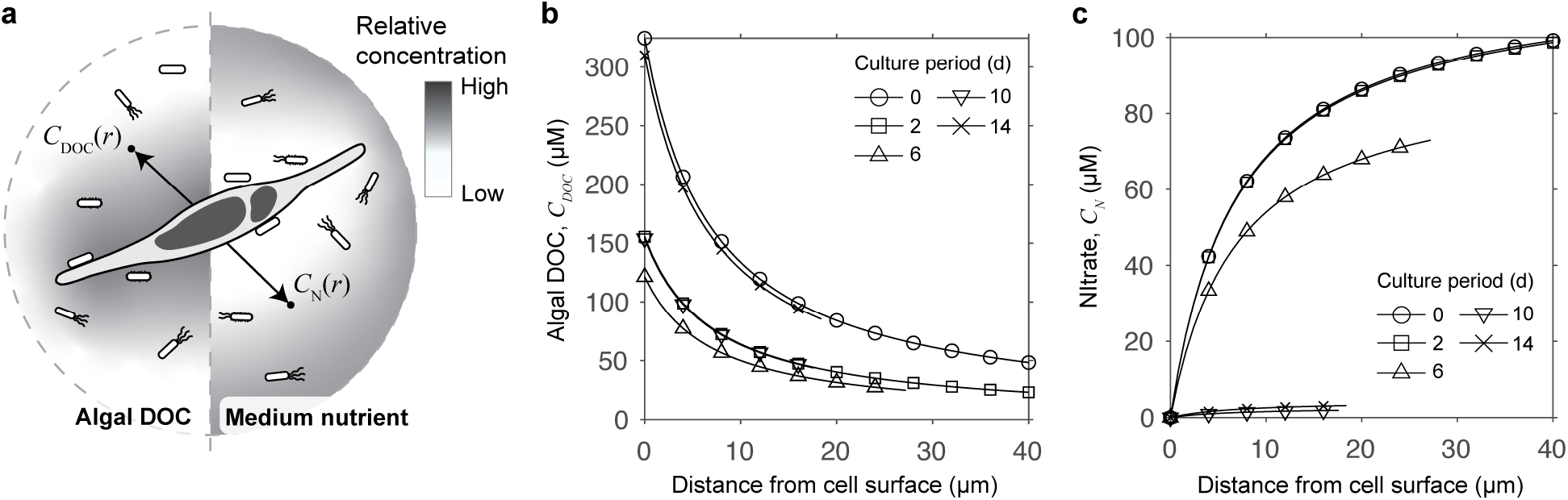
Calculation of nutrient concentrations around an alga. (a) Schematic of two contrasting concentration gradients of algal exudate and medium nutrient at the scale of an alga. Numerical calculation of (b) algal dissolved organic carbon (DOC) and (c) nitrate concentrations measured for 14 days of incubation. in

## Discussion

One major unanswered question in the study of algal-bacterial interactions is how the spatial and temporal dynamics of algal exudation and inorganic nutrient uptake contribute to microbial community development around the phycosphere [10, 49]. Designing experiments to answer such questions with standard laboratory cultures has been particularly difficult, as no method exists to isolate the contributions of these components. We achieved this by using spatial arrays of culture wells with a novel porous microplate and studied how bacteria grew under different nutritional levels of algal DOC and medium-derived inorganic compounds. After analyzing bacterial responses both as single isolates and at the mixed community level, we found evidence that certain taxa became more abundant towards algal-derived organic matter, while others responded more strongly to inorganic nutrients away from the algal cells. The latter finding is unexpected, particularly since the initial bacterial communities originated from algal co-cultures where the only source of organic carbon was algal photosynthesis.

Microalgae are the major consumers of inorganic nutrients in these phototrophic systems, and thus the impact of inorganic nutrients on heterotrophic bacteria in algal co-cultures has remained mostly unexplored, despite the known ability of bacteria to consume inorganic compounds such as nitrate [48, 50–53], phosphate [50, 54] and ammonium [54]. Our co-culture system enabled us to address this underexplored question, and our results further illuminate how bacterial responses are governed by algal culture growth phase and host-bacterial distance (i.e. time and space, respectively). For instance, in early algal growth phase, we observed positive *Marinobacter* population responses towards inorganic nutrients even when the algal host was present (Fig. 3b). However, in later algal growth phase, *Marinobacter* abundances strongly increased with spatial proximity to *P. tricornutum*, likely due to the less diffusible, larger DOC molecules only available in the wells closest to the algae (Fig. 3c).

These experimental observations in the porous microplate enabled us to conceptualize a unique nutrient profile spatially determined by algal growth phase, and we were able to construct a mathematical model of nutrient concentrations around an alga (Fig. 5b), which simultaneously addresses gradients of DOC and inorganic nutrients under different laboratory growth phases. This model allows us to estimate how the algal microbiome might respond to the spatial distribution of the two nutritional factors (algal DOC and inorganic nutrients), because we expect bacteria to reproduce at faster rates in locations where more nutrients are accessible. Specifically for the exponential algal growth stage, we expect bacteria to exhibit a faster growth rate outside of the phycosphere (i.e. free-living), because there are still inorganic nutrients available in the bulk medium and smaller DOC molecules exuded from the algal cells can diffuse further away. On the other hand, for algal cultures in stationary phase, bacteria tend to attach to algal cells or stay within the phycosphere, since inorganic nutrients are depleted throughout the medium and larger DOC molecules diffuse slowly from the surface of the algal cells. Indeed, this scenario explains some experimental observations where the number of attached bacteria increased as algal cultures aged [12, 59].

Notably, by adopting the porous microplate system we constructed a unique environment for algal cells distinct from standard batch cultures, where in the microplate nutrients constantly diffused into the growing culture from outside through the nanoporous HEMA–EDMA. A supply of fresh nutrients into an algal culture is often found in (semi-) continuous systems; however, our system is also distinct from such methods since algae are not removed from the culture and they accumulate to a high cell density. Indeed, *P. tricornutum* reached abundances significantly higher than previously reported values, including under (semi-) continuous systems [60, 61]. We believe this is the first demonstration that algae can accumulate to such high densities, purely driven by a novel cultivation technique. If appropriately scaled up, this finding may provide a substantial impact on developing method for increased biomass production.

Overall, the combination of our experimental culture system and a simple microscale diffusion model successfully explained the temporal and spatial association patterns between algae and different algal-associated bacteria, providing a mechanistic understanding of the distinct impact of inorganic nutrients as well as algal DOC on bacterial growth. We expect this approach can be used in future studies to investigate longstanding questions about community level interactions mediated by metabolite exchange at different spatial scales, including testing hypotheses about bacteria-phycosphere interactions [10, 12, 13], but also more general microbiome interactions in soil, plant, and animal models.

## Acknowledgements

We thank T. J. Samo for isolating *Marinobacter* sp. 3-2, S. Smriga for guidance with culture maintenance, P. Boisvert with SEM imaging, M. Jennings with flow cytometry, S. Kim for advice on statistical analysis. The work was supported by the Department of Energy’s Genome Sciences Program grant SCW1039. Work at LLNL was performed under the auspices of the US Department of Energy at Lawrence Livermore National Laboratory under Contract DE-AC52-07NA27344. H.K. was partly supported by the Kwanjeong Educational Foundation.

## Author contributions

XM and CRB conceived the study. HK and XM designed the experiment. HK and CAV synthesized porous microplates. JRW isolated nucleic acids, prepared libraries and collected sequencing data. JAK and HK performed bioinformatic analysis. HK developed the mathematical model. HK, JAK and XM interpreted the data. All authors wrote the manuscript.

