## Supplementary Information for "Bacterial response to spatial gradients of algal-derived nutrients in a porous microplate"

List of Supplementary Information

- Supplementary Figure S1. Fabrication procedure and dimension of porous microplate.
- Supplementary Figure S2. Growth of *P. tricornutum* in porous microplate-spent medium.
- Supplementary Figure S3. Growth of *Marinobacter* sp. 3-2 and *Algoriphagus* sp. ARW1R1 in the porous microplate.
- Supplementary Figure S4. Cumulative proportion of total variance explained by number of dimensions in principal coordinate analysis.
- Supplementary Table S1. Dehydration measurement of porous microplate well cultures.
- Supplementary Table S2. List of bacterial isolates and their accession numbers.
- Supplementary Table S3. Summary of 16S rRNA sequencing result.
- Supplementary Note S1. Numerical derivation of algal dissolved organic carbon (DOC) and medium nitrate concentrations.

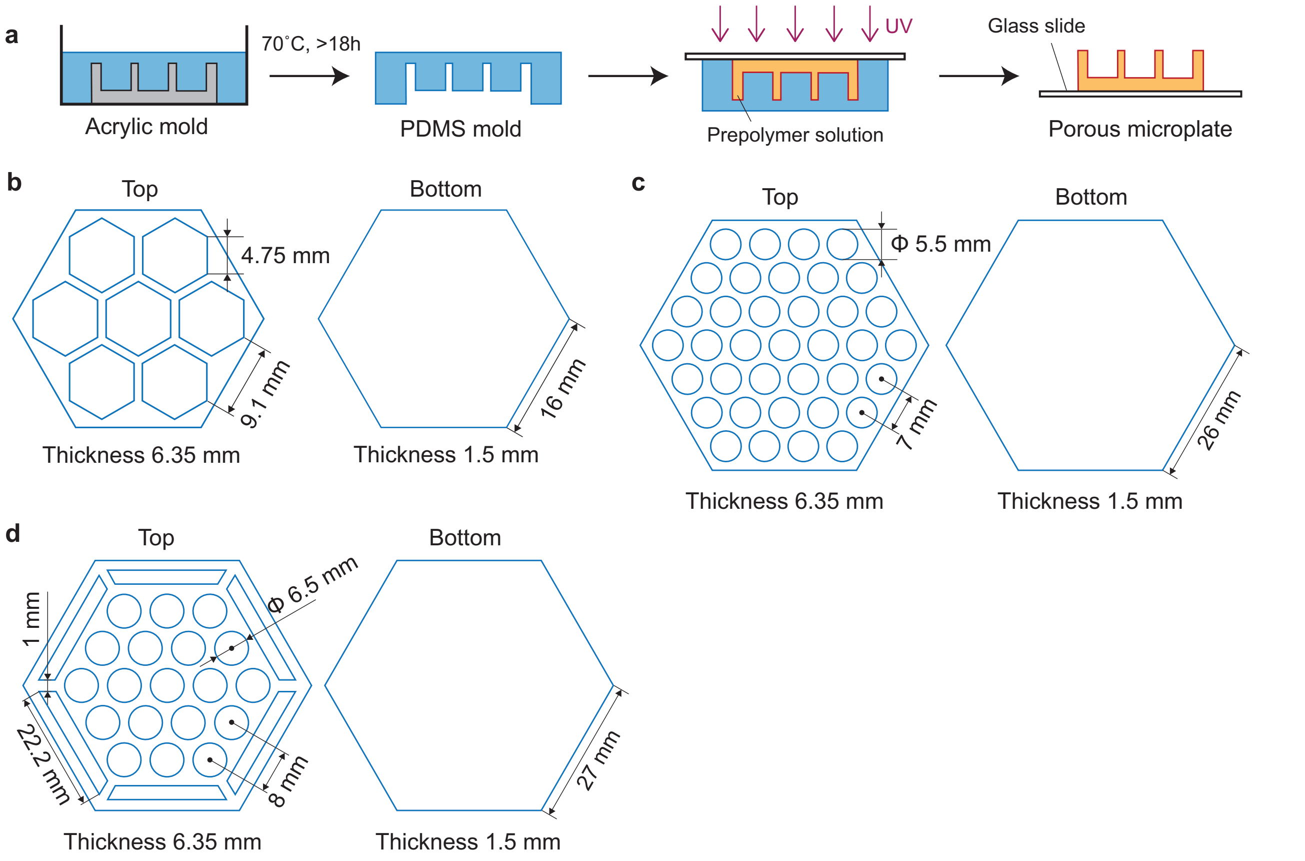


**Supplementary Figure S1. Fabrication procedure and dimension of porous microplate.** (a) Procedure to build a device using PDMS and acrylic molds. Dimension with top and bottom views of a device for (b) *P. tricornutum* incubation experiment, (c) single bacterial isolates growth experiment, and (d) bacterial community analysis experiment.

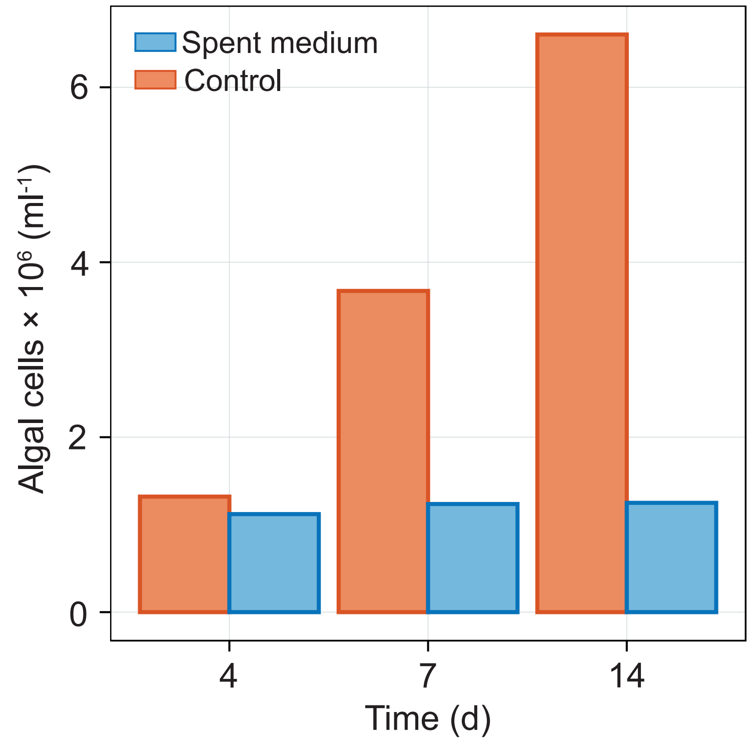


**Supplementary Figure 2. Growth of *P. tricornutum* in porous microplate-spent medium.** Cell numbers of *P. tricornutum* grown in spent f/2-Si medium after the porous microplate incubation experiment (spent) compared to the unaltered medium (control).

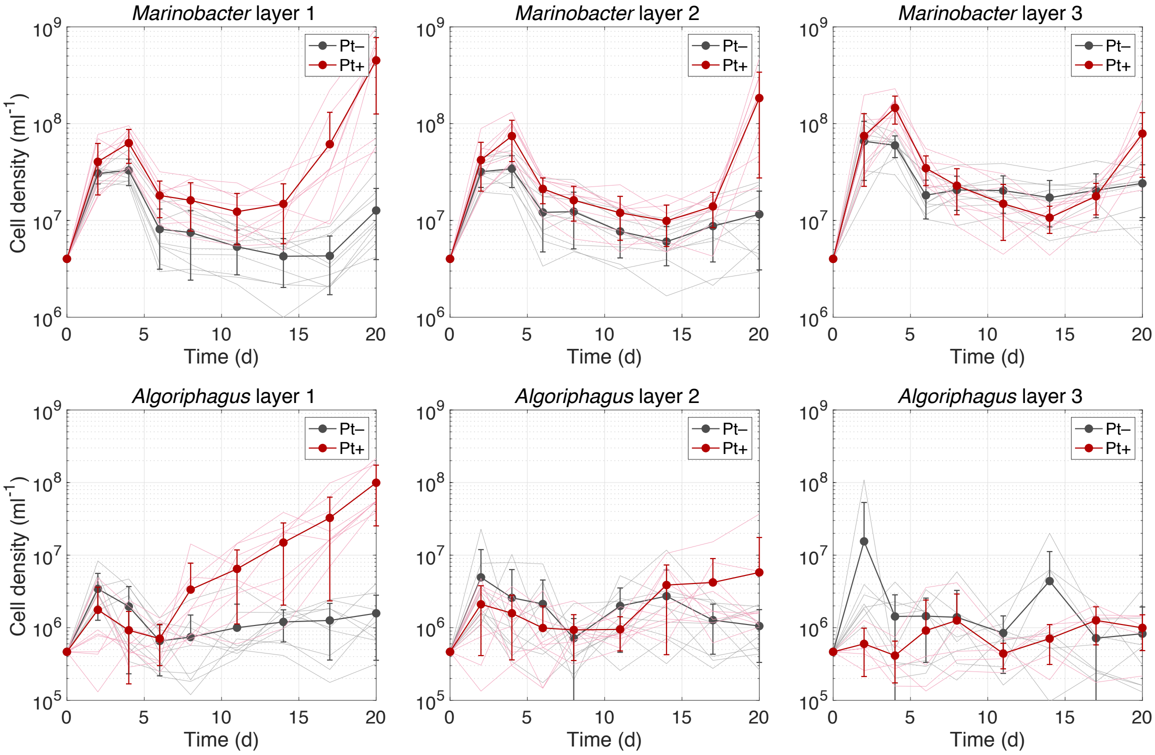


**Supplementary Figure 3. Growth of *Marinobacter* sp. 3-2 and *Algoriphagus* sp. ARW1R1 in the porous microplate.** Bacterial cell densities are plotted by sampling timepoints where they were grown with *P. tricornutum* (red) or without (grey). Error bars, standard deviation of $n = 9$ replicates, except for the following: *Algoriphagus* layer 3 with *P. tricornutum* ($n = 5$), *Algoriphagus* layer 3 without ($n = 7$).

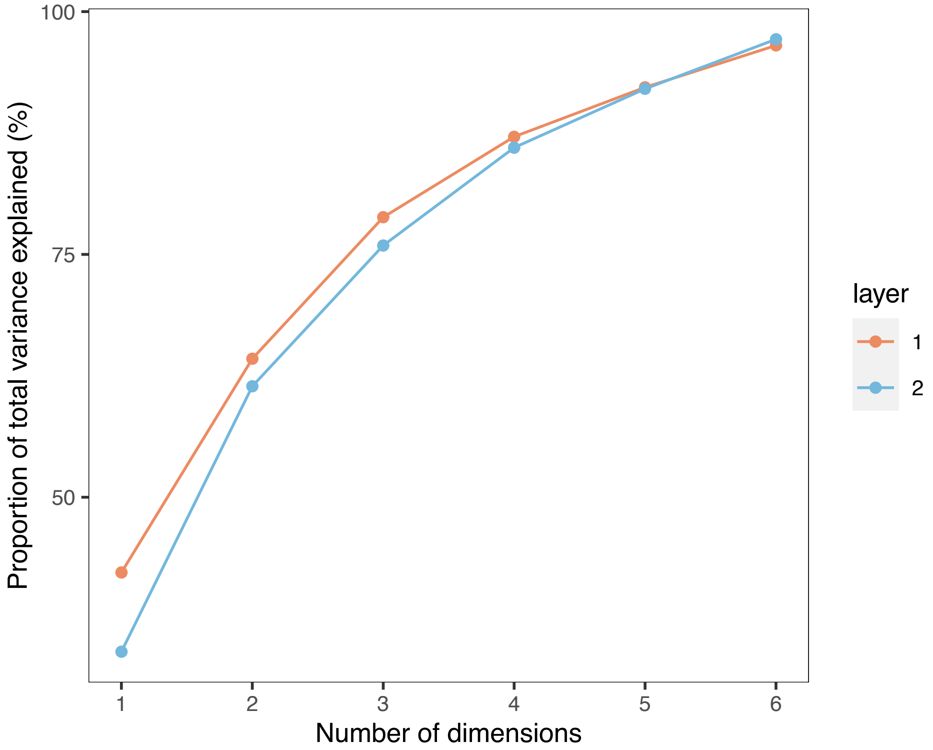


**Supplementary Figure 4. Cumulative proportion of total variance explained by number of dimensions in principal coordinate analysis.**

**Supplementary Table 1. Dehydration measurement of porous microplate well cultures.** Initial *P. tricornutum* culture of 100 µl was inoculated to the hexagonal array of porous microplate then sampled to measure the volume after two weeks of incubation (all units in microliter).

| Location | Replicate 1 | Replicate 2 | Replicate 3 |
|---|---|---|---|
| Center | 90 | 95 | 95 |
| Surrounding 1 | 90 | 100 | 90 |
| Surrounding 2 | 100 | 95 | 90 |
| Surrounding 3 | 100 | 100 | 95 |
| Surrounding 4 | 100 | 90 | 90 |
| Surrounding 5 | 90 | 90 | 95 |
| Surrounding 6 | 100 | 95 | 95 |
| **Total mean** | **94.52** | | |

**Supplementary Table 2. List of bacterial isolates and their accession numbers.**

| Genus | Strain | IMG genome ID | Source |
|---|---|---|---|
| *Marinobacter* | 3-2 | 2785510723 | Samo *et al*. [1] |
| *Algoriphagus* | ARW1R1 | 2747842515 | Samo *et al*. [1] |

**Supplementary Table 3. Summary of 16S rRNA sequencing result.**

| Type | Number of samples | Mean reads | Standard deviation | Minimum reads | Maximum reads |
|---|---|---|---|---|---|
| Negative control | 18 | 19,997 | 42,375 | 61 | 159,617 |
| Phycosphere enrichment (Bacteria only) | 54 | 86,467 | 32,934 | 30,487 | 201,701 |
| Phycosphere enrichment + *P. tricronutum* | 54 | 105,074 | 49,143 | 11,535 | 241,656 |

**Supplementary Note 1. Numerical derivation of algal dissolved organic carbon (DOC) and medium nitrate concentrations.**

We hypothesize that the algal growth phase in laboratory batch cultures determines the spatial gradient of algal DOC and inorganic nutrients. As the cells senesce, the average molecular weight of released compounds increases from low [2–5] to high [2, 5–7], meaning DOC compounds diffuse at slower rates as the cultures age. On the other hand, inorganic nutrients are constantly consumed throughout culture growth so the overall amount will decrease accordingly. This allows us to model the spatial concentration of algal DOC and inorganic nutrients at different growth phases at the scale of a single alga. As a proxy for all inorganic nutrients, nitrate was chosen as it is the inorganic nitrogen source used in our experiments and is expected to show similar diffusivities to other medium components (phosphate, trace metals, etc).

We first begin with Fick's second law of diffusion, $\partial C / \partial t = D\nabla^2 C$ , where $C$ is the concentration of a nutrient and $D$ is the diffusion coefficient. Here we exclude the contribution of fluid advection, which is valid for a cell size of ~10 μm including our model organism *P. tricornutum*, resulting in a small Peclet number ranging from 0.01 to 0.1 [2]. For simplification we assume an alga as a sphere of a radius $r_0$, exuding DOC at a rate $Q_{\mathrm{DOC}}$, and consuming nitrate at a rate $Q_{\mathrm{N}}$ (e.g. mol/s). Under a quasi-steady state where the transient term of the diffusion equation is negligible, we write the concentration of DOC and nitrate as

$$C_{\mathrm{DOC}}(r) = \frac{Q_{\mathrm{DOC}}}{4\pi D_{\mathrm{DOC}} r} \ , \qquad \text{(S1)}$$

$$C_{\mathrm{N}}(r) = C_{\mathrm{inf}} - \frac{Q_{\mathrm{N}}}{4\pi D_{\mathrm{N}} r} \ , \qquad \text{(S2)}$$

where $r$ is radius from the cell center, $C_{\mathrm{inf}}$ is nitrate concentration far away from the cell, and $D_{\mathrm{DOC}}$, $D_{\mathrm{N}}$ are diffusion coefficients of DOC and nitrate respectively [2, 8].

In order to compute $C_{\mathrm{DOC}}(r)$ and $C_{\mathrm{N}}(r)$, we consider a range of domain $r$ with algal cells equally distributed with density $\rho$. Because the average distance between cells is $\sim 2(3/4\pi\rho)^{1/3}$, we denote $R$ by half of the distance, $(3/4\pi\rho)^{1/3}$, and let $r$ range from $r_0$ (surface of a cell) to $R$ (equal distance between two adjacent cells). Noting that nitrate has a relatively low molecular weight whereas DOC compounds can depend on an algal growth phase [2–7], the diffusion coefficients are approximated as $D_N = 10^{-9}$ m$^2$/s, $D_{DOC} = 10^{-9} – 10^{-10}$ m$^2$/s (from exponential to stationary phase).

We note that DOC and nitrate concentrations experimentally measured at *i* th sampling timepoint (day), respectively, denoted by $\overline{C^i_{\mathrm{DOC}}}$ and $\overline{C^i_{\mathrm{N}}}$, can represent spatial average concentrations around the cell. Therefore we write $\overline{C^i_{\mathrm{DOC}}} = \oint_V C_{\mathrm{DOC}}(r)\,dV \cdot V^{-1}$ and $\overline{C^i_{\mathrm{N}}} = \oint_V C_{\mathrm{N}}(r)\,dV \cdot V^{-1}$, where *V* is the volume for which the concentration is spatially averaged within the domain $r \in (r_0, R)$. Following Equations (S1,S2), this allows us to write

$$\overline{C^i_{\mathrm{DOC}}} = \frac{3Q^i_{\mathrm{DOC}}}{8\pi D_{DOC}} \cdot \frac{R^2 - r_0^2}{R^3 - r_0^3}, \tag{S3}$$

$$\overline{C^i_{\mathrm{N}}} = C^i_{\mathrm{inf}} - \frac{3Q^i_{\mathrm{N}}}{8\pi D_{\mathrm{N}}} \cdot \frac{R^2 - r_0^2}{R^3 - r_0^3}, \tag{S4}$$

where physical quantities with upper index *i* denote those measured (or computed) on *i* th sampling day.

Additionally, we assume at the surface of a cell that nitrate concentration is nearly zero as the nutrients are quickly consumed by algae, which has experimentally been shown with nitrate and *P. tricornutum* [9]. This allows us to construct a boundary condition at $r = r_0$ of the nitrate concentration, giving

$$0 = C^i_{\mathrm{inf}} - \frac{Q^i_{\mathrm{N}}}{4\pi D_{\mathrm{N}} r_0} . \tag{S5}$$

We estimated values of $Q^i_{\mathrm{DOC}}$, $Q^i_{\mathrm{N}}$, $Q^i_{\mathrm{inf}}$ by solving Equations (S3–S5) and obtained nutrient concentration profiles with Equations (S1,S2). Among the four model species reported by Biddanda *et al.* [5], the concentrations and abundance data were chosen from the cyanobacterium *Synechococcus bacillaris*, because its cell abundances were the most similar to that of *P. tricornutum*, giving an appropriate range of distance domain *r*.

Numerical calculations for Figure 4b were performed with growth data of *S. bacillaris* on day 0–14 by using MATLAB (Mathworks).
